# The Pax9/Wnt pathway regulates secondary palate formation in mice

**DOI:** 10.1101/110692

**Authors:** Shihai Jia, Jing Zhou, Christopher Fanelli, Yinshen Wee, John Bonds, Pascal Schneider, Gabriele Mues, Rena N. D’Souza

## Abstract

Clefts of the palate and/or lip arise in about 1/700 human live births and are caused by multiple genetic and environmental factors. Studies of mouse knockout models of cleft palate have improved our understanding of the molecular control of palatogenesis. While it is known that Pax9 regulates palatogenesis through Bmp, Fgf and Shh signaling, there is still much to learn about its precise relationship with other pathways. Here we show that alterations of Wnt expression and decreased Wnt activity in *Pax9^-/-^* palatal shelves are a result of Pax9’s ability to directly bind and repress the promoters of *Dkk1* and *Dkk2,* proteins that antagonize Wnt signaling. The delivery of small-molecule Dkk inhibitors (Wnt agonists) into the tail-veins of pregnant *Pax9^+/-^* mice from E10.5 to E14.5 restored Wnt signaling, promoted cell proliferation, bone formation and restored the fusion of palatal shelves in *Pax9^-/-^* embryos. In contrast, other organ defects in *Pax9* mutants were not corrected. These data uncover a unique molecular relationship between *Pax9* and Wnt genes in palatogenesis and offer a new approach for treating cleft palates in humans.

**Summary Statement:** These studies demonstrate that the Pax9/Wnt genes regulate murine palatogenesis. This unique molecular relationship is proven by the correction of cleft defects in *Pax9-*deficient mice through Wnt agonist therapies.

## Introduction

Development of the mammalian palate involves a complex and tightly coordinated series of events that regulate cell migration, proliferation, growth and differentiation as well as programmed cell death. First, the medial nasal processes fuse with the maxillary prominences to form the upper lip, and then extend orally to give rise to the presumptive primary and secondary palates, respectively. As morphologically distinct structures, the secondary palatal shelves undergo a critical reorientation from a vertical to a horizontal position. Palatal shelf migration toward the midline continues until fusion occurs. Separation of the oral and nasal cavities is evident after complete fusion of the secondary palate with the primary palate and nasal septum is complete (Mossey et al., 2009; Greene and Pisano, 2010; Bush and Jiang, 2012). Failures in palatogenesis occur during the elevation, migration or fusion of palatal shelves and serve as the best evidence that these processes are under strict molecular control. It is hence not surprising that clefts of the secondary palate, that occur singly or in combination with cleft lip, are the most common among human craniofacial birth defects affecting 1 in 700 live births (Dixon et al., 2011; Watkins et al., 2014). Affected individuals suffer difficulties with feeding, speech, hearing, cognition and social integration and must endure surgeries and multidisciplinary care throughout life.

Advances in sequencing and genotyping technologies along with molecular and epidemiological studies have helped unravel the complex etiology of secondary palate defects in humans (Mossey et al., 2009; Dixon et al., 2011). Since morphological events in mice closely resemble that seen in humans, mouse genetic models with secondary palate defects have also been valuable in dissecting the signaling pathways that drive palatogenesis (Greene and Pisano, 2010; Bush and Jiang, 2012; Funato et al., 2015). Taken together, these studies reveal that palatogenesis involves a tight interplay of gene networks and environmental risk factors that regulate a temporospatial series of molecular, cellular and tissue interactions. Gene ontology studies of cleft palate defects in mice reveal that more than half of the genes involved are transcription factors, such as, *Msx1* (Satokata and Maas, 1994; Zhang et al., 2002), *Pax9* (Peters et al., 1998; Zhou et al., 2011), *Osr2* (Lan et al., 2004), *Gbx2* (Byrd and Meyers, 2005), *Tbx22* (Pauws et al., 2009), *Tfap2A* (Brewer et al., 2004), *Hsoxa2* (Gendron-Maguire et al., 1993). Moreover, growth factor families such as TGF-ßs, Shh, Fgf, and Wnt along with membranous molecules and extracellular matrix proteins are also implicated as playing important roles in palate formation (Jiang et al., 2006; Funato et al., 2015).

Among the mouse genetic models of cleft palate that are available, the *Pax9^-/-^* mouse model has provided valuable insights into the mechanisms that lead to the failure of palatogenesis. Pax9 belongs to the family of paired-box DNA-binding domain containing transcription factors (Stapleton et al., 1993). *Pax9^-/-^* mice consistently exhibit clefts of the secondary palate and die shortly after birth. Other phenotypic defects include: agenesis of the thymus, parathyroid glands, ultimobranchial bodies and teeth (Peters et al., 1998; Zhou et al., 2011). Molecular and mouse genetic approaches have demonstrated the importance of *Pax9* in the patterning of the anterior-posterior (AP) axis and outgrowth of palatal shelves through its control of cell proliferation and its interactions with the Bmp, Fgf and Shh pathways (Zhou et al., 2013). However, there is much to learn about Pax9’s relationship with other signaling pathways in this process. For example, the Wnt genes encode a family of secreted signaling proteins that regulate developmental processes that are critical in palate formation, such as early patterning, cell proliferation, differentiation and apoptosis. Yet, little is known about the nature of Pax9’s relationship with Wnt genes interact during palate formation.

In these studies we demonstrate that *Pax9* is upstream of Wnt signaling during palatogenesis. Our gene expression analyses and molecular assays demonstrate that the inhibitors of Wnt expression, namely, *Dkk1* and *Dkk2* directly interact with Pax9 offering an explanation for why these genes are down regulated in *Pax9* mutant palatal mesenchyme. The controlled intravenous delivery of Wnt agonist small molecules to pregnant *Pax9^+/-^* mice consistently corrects cleft palate defects in mutant progeny thus confirming that the Pax9-Wnt signaling pathway is a critical regulator of palatogenesis.

## Results

### Pax9 shares a direct upstream molecular relationship with the Wnt signaling pathway during palatogenesis

We first examined the molecular consequences of *Pax9* deficiency in forming palatal shelves at E13.5 using RNASeq analyses. Our data indicate that the complete lack of *Pax9* led to more than a 1.5-fold (p<0.01) change in expression of approximately 1368 genes among which the Wnt pathway genes appear strongly enriched (Table 1, left column). A key observation is the increased expression of *dickkopf* genes, *Dkk1* and *Dkk2,* which encode extracellular proteins that block the binding of Wnt ligands to the Lrp5/6 receptor complex. Genes involved in Bmp, Eda, Fgf and Shh signaling were also affected (Table1, right column), confirming previous findings (Zhou et al., 2013). Alterations in gene expression levels were confirmed by quantitative real-time PCR analysis (Fig. 1), that showed the expression of Wnt signaling pathway inhibitors *Dkk1* and *Dkk2* had increased 1.7 fold, and the expression of *Msx1, Lef1, Cited1* and *Gbx2* had decreased more than 2 fold in the *Pax9* mutant samples.

**Table 1.** Expression profiles of *Pax9* affected genes with functional enrichment analysis in E13.5 palatal shelves

| Gene | *Pax9*<sup>+/-</sup> | *Pax9*<sup>-/-</sup> | FC | pvalue | Gene | *Pax9*<sup>+/-</sup> | *Pax9*<sup>-/-</sup> | FC | pvalue |
| --- | --- | --- | --- | --- | --- | --- | --- | --- | --- |
| *Dkk1* | 180.64 | 324.53 | 1.80 | 3.88E-03 | *Edar* | 184.38 | 98.87 | -1.86 | 3.41E-05 |
| *Dkk2* | 3199.37 | 4888.36 | 1.52 | 8.71E-13 | *Tfap2b* | 2405.06 | 873.21 | -2.75 | 3.22E-85 |
| *Wnt7a* | 25.82 | 6.91 | -3.74 | 0.015 | *Msx1* | 6554.76 | 3221.79 | -2.03 | 2.22E-41 |
| *Wnt3* | 42.53 | 20.30 | -2.09 | 5.58E-03 | *Msx2* | 477.41 | 265.37 | -1.80 | 4.58E-04 |
| *Wnt9b* | 65.42 | 35.74 | -1.83 | 3.34E-03 | *Osr2* | 6338.19 | 11254.15 | -1.72 | 7.98E-24 |
| *Lef1* | 1231.91 | 600.32 | -2.05 | 1.02E-16 | *Foxf1* | 5657.10 | 3784.74 | -1.50 | 3.81E-10 |
| *Rspo1* | 982.53 | 631.61 | -1.56 | 8.00E-03 | *Hand2* | 165.14 | 63.62 | -2.59 | 1.06E-10 |
| *Gbx2* | 26.34 | 6.24 | -4.22 | 4.29E-05 | *Cited1* | 404.22 | 165.68 | -2.44 | 1.28E-07 |
| *Gad1* | 32.42 | 4.53 | -7.16 | 6.87E-14 | *Ret* | 550.22 | 149.65 | -3.68 | 1.99E-22 |
| *Tbx20* | 30.41 | 11.31 | -2.69 | 0.016 | *Hoxa2* | 28.37 | 2.77 | -10.24 | 3.39E-03 |
| *Fgf4* | 109.69 | 9.22 | -11.90 | 2.93E-05 | *Ryr1* | 876.49 | 2062.90 | 2.35 | 7.30E-03 |

The expression levels were shown by normalized counts. FC, Fold Change; -, down-regulation.

**Figure 1.**
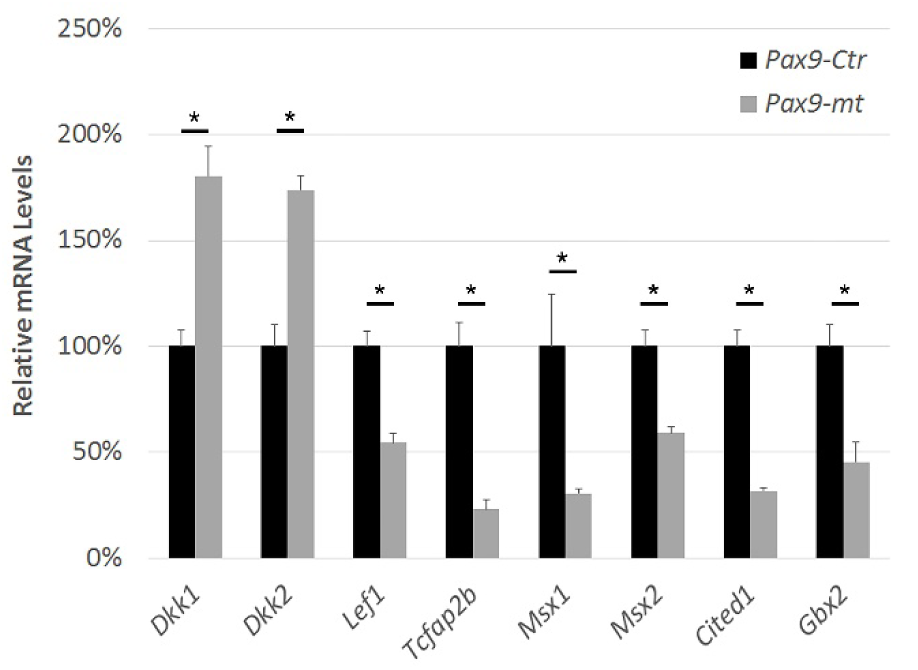
The differentially expressed genes in E13.5 *Pax9^-/-^* palatal shelves were confirmed by real-time PCR. The mRNA levels of selected differentially expressed genes were verified by RT-qPCR (n=5). Error bars indicate SEM, * *P* < 0.01.

The relationship between Pax9 and the Wnt pathway was further assessed by studying whether the expression of *Dkk1* and *Dkk2* overlapped with that of *Pax9* during normal palatogenesis. As shown in Fig. 2A-F, *Dkk1* and *Dkk2* expression levels increase from E13.5 to E14.5 and appears restricted to posterior palatal mesenchyme where *Pax9* is also dominantly expressed (Fig. 2A-F). Interestingly, these overlapping domains of *Dkk1, Dkk2* and *Pax9* expression in posterior palatal mesenchyme differ from that of *Msx1* and *Bmp4* which localize to the anterior region of the palatal mesenchyme (Zhang et al., 2002; Han et al., 2009; Fuchs et al., 2010). In situ hybridizations showed that expression of both Wnt inhibitors increased in the absence of *Pax9* palate mesenchyme (Fig. 2G-N). At E13.5, *Dkk1* expression was detected in the posterior palatal mesenchyme with a buccal to lingual gradient pattern (Fig. 2H). In *Pax9* mutant samples, *Dkk1* expression was increased in the posterior palate and extended to anterior region (Fig. 2I,J). *Dkk2* expression was higher in the posterior palatal mesenchyme with lingual to buccal gradient (Fig. 2L), and increased in the *Pax9* mutant samples (Fig 2N).

**Figure 2.**
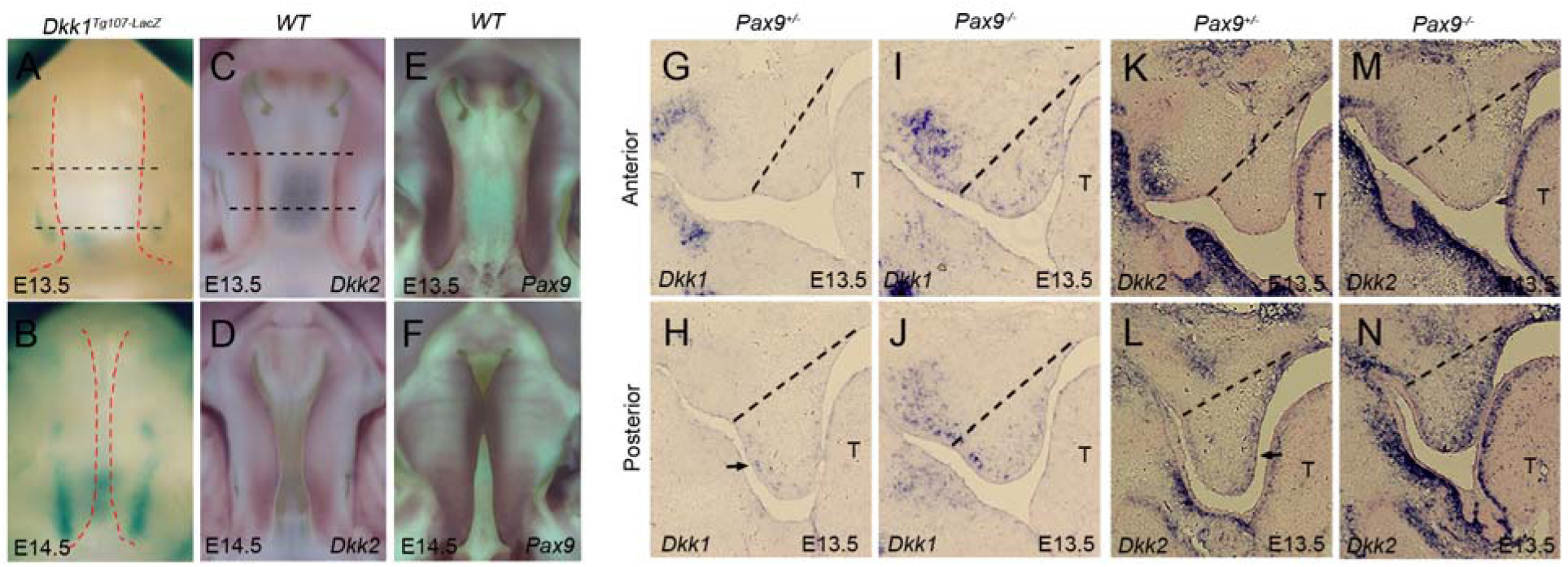
The expression of *Pax9* overlapped and modulated the expression of *Dkk1* and *Dkk2* in developing palate. The whole mount view of expression patterns of Dkk1, Dkk2 and Pax9 in developing palate (A-F). The whole mount LacZ staining of developing palate shelves in *Dkk1^Tg107-LacZ^* embryos at E13.5 (A) and E14.5 (B), red dashed lines indicate the boundary of palate, black dashed lines indicate the position of the sections in the G,H. The whole mount *In-situ* hybridization of *Dkk2* (purple) in wildtype embryos at E13.5 (C) and E14.5 (D), black dashed lines indicate the position of the sections in the K,L. The whole mount *In-situ* hybridization of *Pax9* (purple) in wildtype embryos at E13.5 (E) and E14.5 (F). The expressions of *Dkk1* and *Dkk2* were increased in *Pax9^-/-^* palate (G-N). *In-situ* hybridization of *Dkk1* mRNA (blue) in sections of anterior (G,I) and posterior (H,J) palate from *Pax9^+/-^*and *Pax9^-/-^.* T, tongue; dashed lines indicate the boundary of palate, black arrow indicates the buccal side *Dkk1 in-situ* signals. *In-situ* hybridization of *Dkk2* mRNA (blue) in sections of anterior (K,M) and posterior (L,N) palate from *Pax9^+/-^* and *Pax9^-/-^.* T, tongue; dashed lines indicate the boundary of palate, black arrow indicates the lingual side *Dkk2 in-situ* signals. n=6 for whole mount in-situ hybridization, n=5 for sectioned in-situ hybridization.

In order to elucidate the nature of the putative upstream relationship of Pax9 to the Wnt inhibitors, we performed ChIP-qPCR analysis. Fig. 3B,C shows that Pax9 directly interacts with promoters of *Dkk1* and *Dkk2* and that its binding was enriched 3- to 6-fold when compared to negative controls (Fig. 3B,C). Taken together, these data suggest for the first time that Pax9 shares a direct upstream molecular relationship with Wnt inhibitor genes during key stages in palatogenesis.

**Figure 3.**
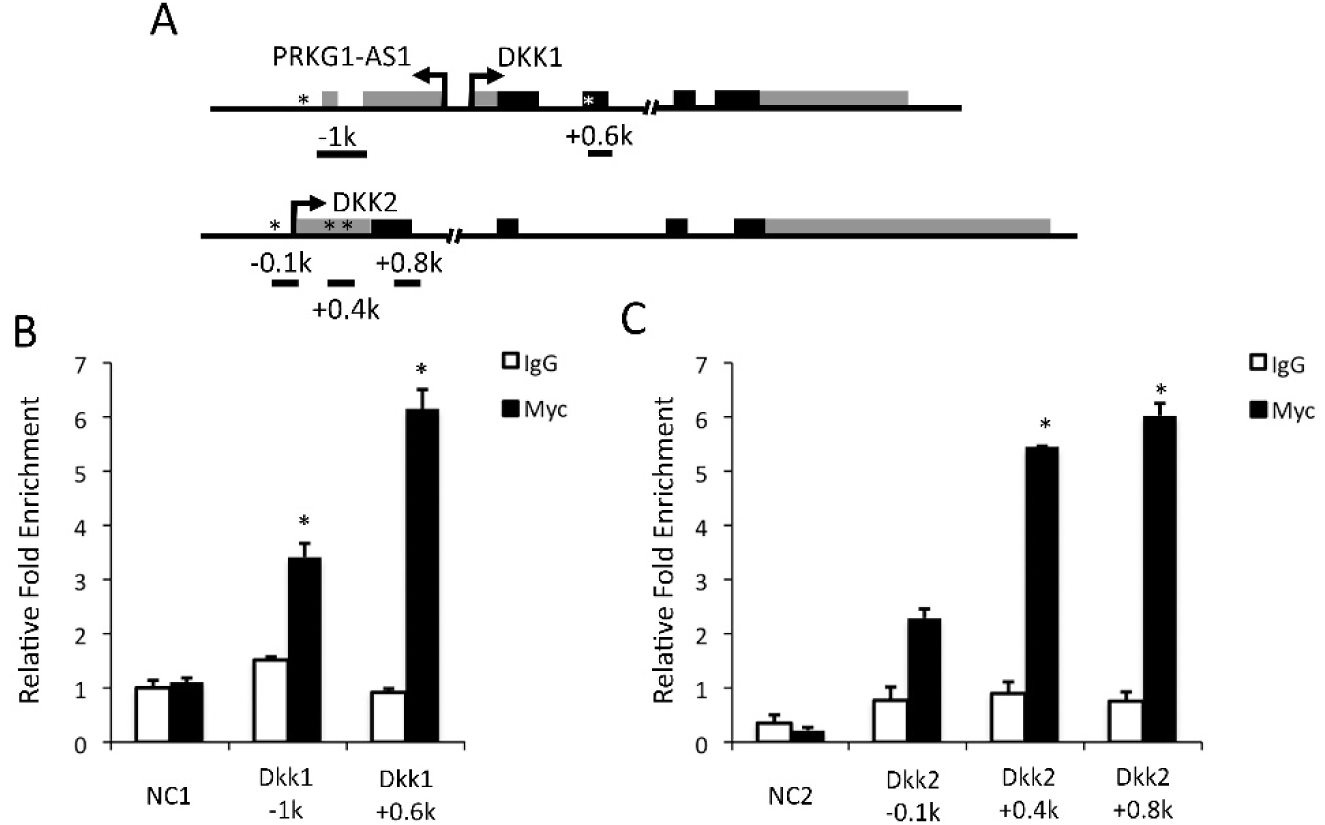
Pax9 had binding activity on the promoters of *Dkk1* and *Dkk2.* (A) Schematic of promoter region of *Dkk1* and *Dkk2,* * indicates the potential binding site. (B) The enrichment of Pax9 binding fragments in *Dkk1* promoter region were verified by qPCR (n=5). Error bars indicate SEM, * *P* < 0.01. NC1, non-binding control region 1. (C) The enrichment of Pax9 binding fragments in *Dkk2* promoter region were verified by qPCR (n=5). Error bars indicate SEM, * *P* < 0.01. NC2, non-binding control region 2.

### Small molecule Wnt agonists that act as Dkk inhibitors cure cleft palate defects in *Pax9^-/-^* mice confirming that the Pax9/Wnt pathway is critical in palatogenesis

We assessed whether the intravenous (IV) administration of small-molecules that exclusively target the Wnt pathway can correct palatal defects that are a consistent phenotype in *Pax9^-/-^* mice. The small-molecule WAY-262611, that potentiates Wnt-β- catenin signaling by inhibiting Dkk1 (Pelletier et al., 2009) was administered daily at 12.5 mg/kg and 25 mg/kg into the tail veins of pregnant *Pax9^+/-^* mice from E10.5 to E14.5 covering the palatal stages of initiation, outgrowth and elevation (Fig. 4). The delivery of 5 doses of WAY-262611 (Fig. S1A) at either level resulted in fusion of the secondary palate in all (18/18) *Pax9^-/-^* pups examined. Treatment from E10.5 to E13.5 rescued the cleft phenotype, but to a lesser extent than the E10.5 to E14.5 regimen (Table. 2). As disclosed in Table 2, 60% (11/18) additionally displayed complete fusion between the primary and secondary palates while 7/18 showed small residual defects in the zone of fusion of the primary and secondary palates around the 3^rd^ rugae (Fig. S2B). Since *Pax9^-/-^* pups die postnatally, E18.5 palatal shelves with small-sized fusion defects after treatment with WAY-262611 were cultured: 3 days later, all palates displayed complete fusion (Fig. S2). This suggests that the lack of complete fusion was due to a timing delay rather than to genetic differences between the primary and secondary palates.

**Figure 4.**
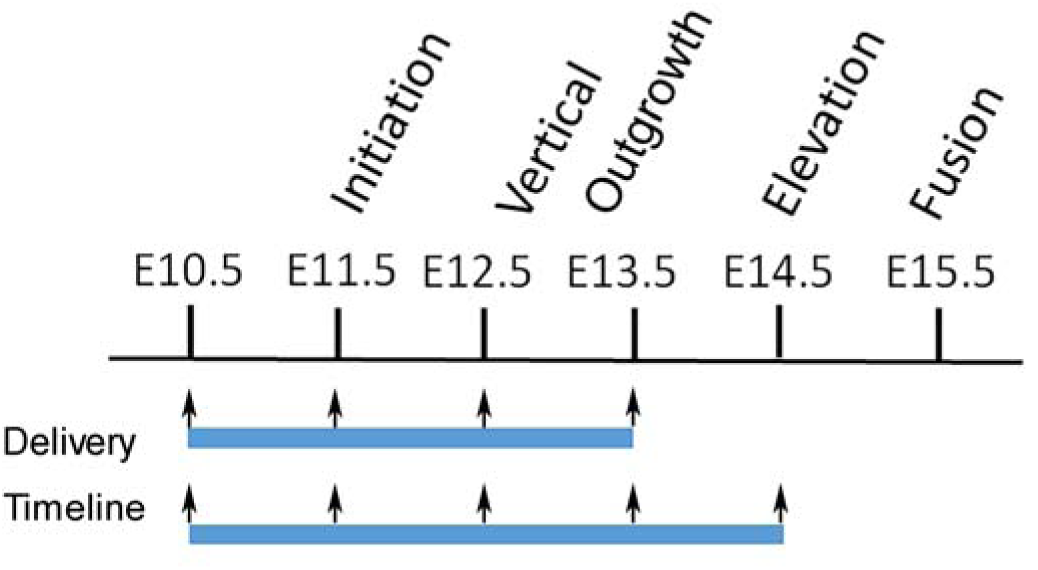
Experimental design. WAY-262611 or IIIc3a was intravenously injected through tail vein during the designed periods (blue bars). Black arrows indicate daily injection.

**Table 2.** Secondary palate closure with treatments of Wnt agonists.

| Drugs | Expt. Groups | Time Line | Dosage (mg/Kg) | *Pax9*<sup>-/-</sup> No. | Closure | Cleft Palate |
| --- | --- | --- | --- | --- | --- | --- |
| WAY-262611 | 1 | E10.5-14.5 | 12.5 | 6 | 4/2* | 0 |
|  | 2 | E10.5-14.5 | 25 | 12 | 7/5* | 0 |
|  | 3 | E10.5-13.5 | 12.5 | 8 | 2/1* | 5 |
|  | 4 | E10.5-13.5 | 25 | 9 | 3/3* | 3 |
| Illc3a | 5 | E10.5-14.5 | 12.5 | 9 | 4/3* | 2 |
|  | 6 | E10.5-14.5 | 25 | 6 | 2/3* | 1 |

\*, number of *Pax9*<sup>-/-</sup> embryos with complete fusion between the primary and secondary palates.

We tested another small-molecule agonist of Wnt signaling, IIIc3a, that blocks the Wnt antagonists Dkk1, 2 and 4 (Li et al., 2012) (Fig. S1B). Daily IV delivery of IIIc3a to pregnant dams from E10.5 to E14.5 resulted in secondary palate fusion in 80% (12/15) of *Pax9^-/-^* pups and palate closure in 40% (6/15). Hematoxylin & eosin as well as Masson’s trichrome stains revealed that both pharmacological interventions brought about a true correction of the secondary palatal defect and the deposition of collagen-enriched matrix in the area of the midline (Fig. 5). The sample size for each treatment group were confirmed by power analysis using PASS software, version 11 (Hintze, J. (2011). PASS 11. NCSS, LLC. Kaysville, Utah, USA) provided by University of Utah Study Design and Biostatistics Center to reach power > 80%.

**Figure 5.**
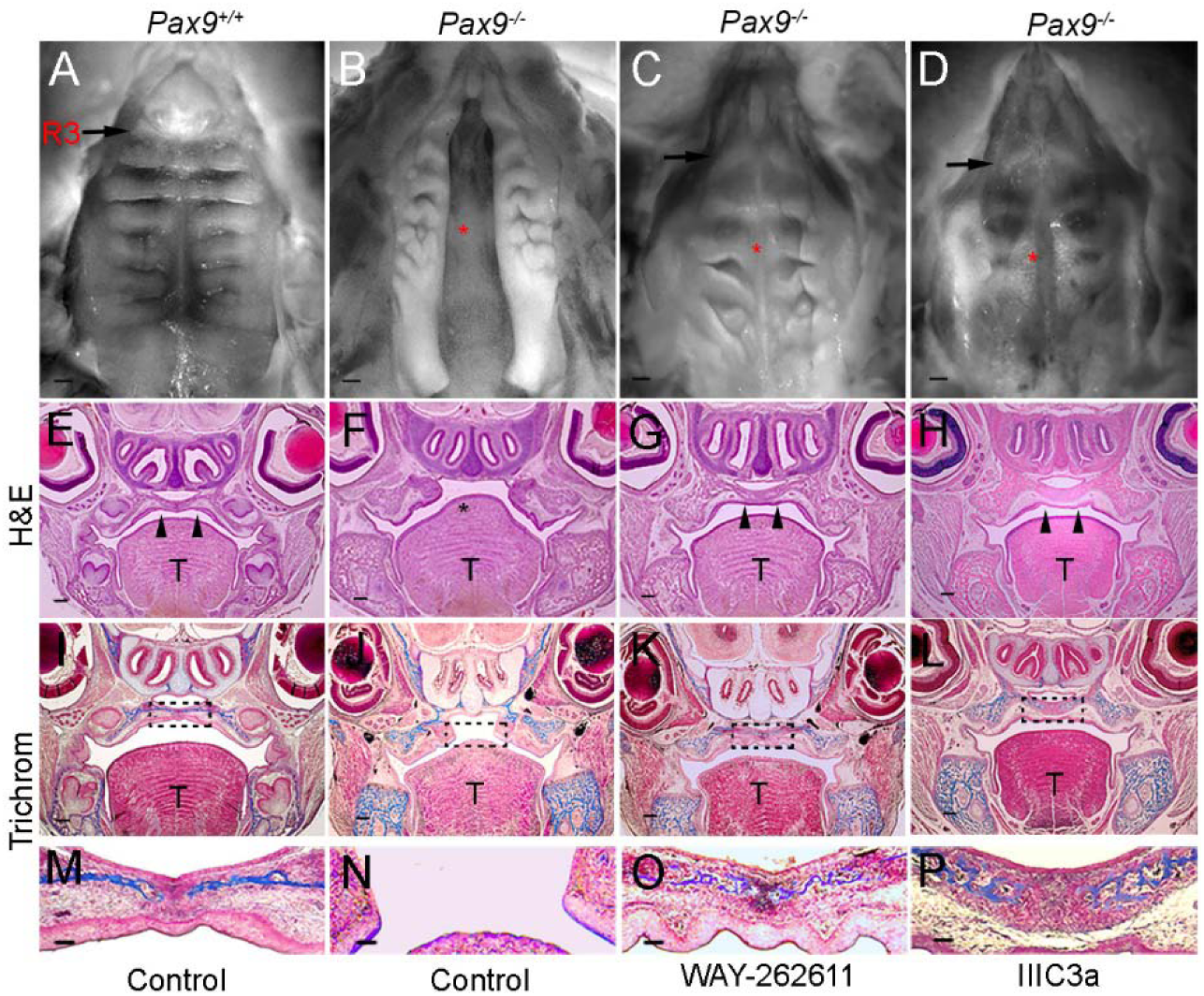
Secondary palate fusion in *Pax9^-/-^*embryos after treatments. Representative photographs of palate formation in control treatment *Pax9^+/+^* (A,E,I,M), *Pax9^-/-^*(B,F,J,N), WAY-262611 treated *Pax9^-/-^* (C,G,K,O) and IIIc3a treated *Pax9^-/-^* (D,H,L,P) at E18.5. The whole mount oral view of palatal shelf showed cleft palate in control treatment *Pax9^-/-^* embryos (n=10) (B) and intact palate with disordered rugaes in WAY-262611 (n=18) or IIIc3a (n=12) treated *Pax9^-/-^* embryos (C,D). R3, Rugae 3; Black arrows indicate the 3^rd^ rugae; red * indicates cleft palate or secondary palate fusion after treatment; HE staining (E-H) and Trichrom staining (I-P) of frontal sections showed that the palatal shelves were fused in control treatment Pax9^+/+^(E,I,M), WAY-262611 treated *Pax9^-/-^* (G,K,O) and IIIc3a treated *Pax9^-/-^* (H,L,P). T, tongue; arrow heads indicate intact palate shelve; dashed rectangle in (I-L) represents the enlarged area of (M-P).

Since Pax9 regulates *Msx1,* a transcription factor that also plays a role in palatogenesis, IV injections of WAY-262611 and III3ca at 25 mg/kg were given to pregnant *Msx1^+/-^* mice, but this failed to rescue cleft palate defects in mutant pups (0/50) (Fig. S3). Since *Msx1* expression is spatially restricted to the anterior region of palatal mesenchyme where *Dkk1* and *Dkk2* expression levels are reduced, it is likely that the reactivation of Wnt signaling only influenced downstream events in posterior regions of the palate where *Pax9* is more dominantly expressed.

### The *in-utero* delivery of small molecule Wnt agonists activated Wnt activity that led to an increase in cell proliferation, growth and fusion of the palatal shelves

Evidence that these pharmacological interventions activated Wnt signaling in palatal mesenchyme comes from the increased levels of *β-catenin* expression that became visible in *Pax9^-/-^* palatal mesenchyme after treatment with WAY-262611 or IIIc3a (Fig. 6A). RNASeq data showed increased levels of Wnt target gene expression in treated palates (Table 3). WAY-262611 had no effect on *Dkk1* and *Dkk2* mRNA levels, because it acts at the protein level. As an indicator of restored Wnt signaling after treatment, real-time PCR result shows that the 6-fold level in reduction of *Gbx2 and L1cam,* which are direct Wnt target genes, were partially rescued (Fig. 6B). *Gbx2* mutant mice have cleft palate defects while L1cam-deficient mice show other craniofacial defects (Byrd and Meyers, 2005).

**Figure 6.**
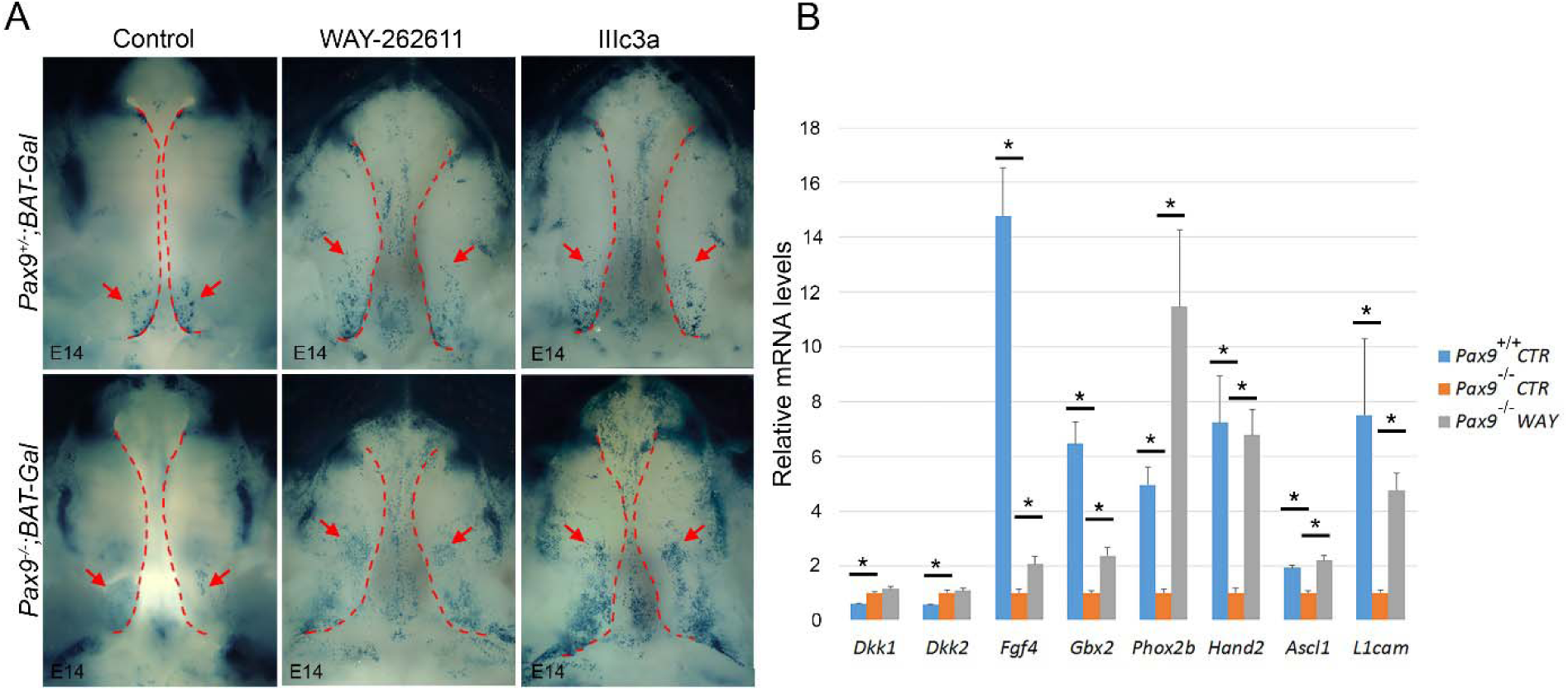
Wnt signaling activity and the expression of downstream targets were partially restored in *Pax9*^-/-^ palate after treatments. (A) The whole mount LacZ staining (blue) at E14.0 showed that the Wnt signaling activity was reduced in the posterior palate of *Pax9^-/-^;BAT-Gal* mouse compared with *Pax9^+/-^;BAT-Gal,* after WAY-262611 or IIIc3a treatment, the Wnt signaling activities were restored in the palate of *Pax9^-/-^;BAT-Gal* mouse. Red arrows point the Wnt signaling activity in palate; dashed line indicates the boundary of palate. (B) The mRNA levels of selected differentially expressed genes were verified by RT-qPCR (n=5). Error bars indicate SEM, * *P* < 0.01.

**Table 3.**
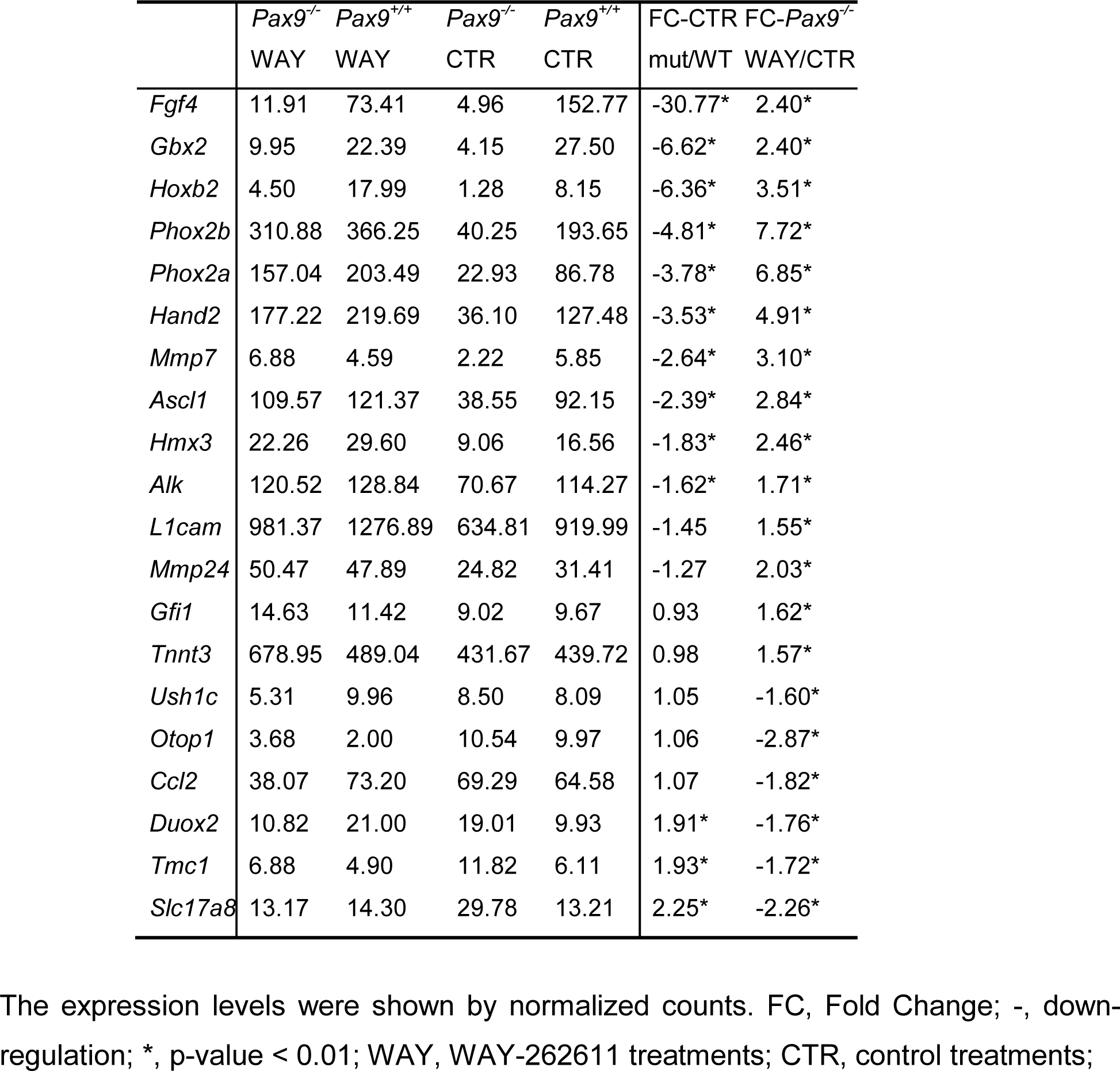
Differentially Expressed Genes in *Pax9^-/-^* Samples with or without Treatment

|  | *Pax9*<sup>-/-</sup><br>WAY | *Pax9*<sup>+/+</sup><br>WAY | *Pax9*<sup>-/-</sup><br>CTR | *Pax9*<sup>+/+</sup><br>CTR | FC-CTR<br>mut/WT | FC-*Pax9*<sup>-/-</sup><br>WAY/CTR |
| --- | --- | --- | --- | --- | --- | --- |
| *Fgf4* | 11.91 | 73.41 | 4.96 | 152.77 | -30.77* | 2.40* |
| *Gbx2* | 9.95 | 22.39 | 4.15 | 27.50 | -6.62* | 2.40* |
| *Hoxb2* | 4.50 | 17.99 | 1.28 | 8.15 | -6.36* | 3.51* |
| *Phox2b* | 310.88 | 366.25 | 40.25 | 193.65 | -4.81* | 7.72* |
| *Phox2a* | 157.04 | 203.49 | 22.93 | 86.78 | -3.78* | 6.85* |
| *Hand2* | 177.22 | 219.69 | 36.10 | 127.48 | -3.53* | 4.91* |
| *Mmp7* | 6.88 | 4.59 | 2.22 | 5.85 | -2.64* | 3.10* |
| *Ascl1* | 109.57 | 121.37 | 38.55 | 92.15 | -2.39* | 2.84* |
| *Hmx3* | 22.26 | 29.60 | 9.06 | 16.56 | -1.83* | 2.46* |
| *Alk* | 120.52 | 128.84 | 70.67 | 114.27 | -1.62* | 1.71* |
| *L1cam* | 981.37 | 1276.89 | 634.81 | 919.99 | -1.45 | 1.55* |
| *Mmp24* | 50.47 | 47.89 | 24.82 | 31.41 | -1.27 | 2.03* |
| *Gfi1* | 14.63 | 11.42 | 9.02 | 9.67 | 0.93 | 1.62* |
| *Tnnt3* | 678.95 | 489.04 | 431.67 | 439.72 | 0.98 | 1.57* |
| *Ush1c* | 5.31 | 9.96 | 8.50 | 8.09 | 1.05 | -1.60* |
| *Otop1* | 3.68 | 2.00 | 10.54 | 9.97 | 1.06 | -2.87* |
| *Ccl2* | 38.07 | 73.20 | 69.29 | 64.58 | 1.07 | -1.82* |
| *Duox2* | 10.82 | 21.00 | 19.01 | 9.93 | 1.91* | -1.76* |
| *Tmc1* | 6.88 | 4.90 | 11.82 | 6.11 | 1.93* | -1.72* |
| *Slc17a8* | 13.17 | 14.30 | 29.78 | 13.21 | 2.25* | -2.26* |

4 The expression levels were shown by normalized counts. FC, Fold Change; -, down- 5 regulation; \*, p-value < 0.01; WAY, WAY-262611 treatments; CTR, control treatments;

*Pax9* deficiency lowers cell proliferation in palatal mesenchyme as reported by (Zhou et al., 2013), the posterior palate region had significant reduction in BrdU incorporation. After treatment with Wnt signaling agonist, the cell proliferation was restored to a level that appeared adequate for the outgrowth of palatal shelves towards the midline (Fig. 7A,B). The BrdU labeling assay in posterior palate at E14.0 showed that the ratios of BrdU positive palatal mesenchymal cells were significantly reduced in the *Pax9* mutant samples confirming previous reports (Zhou et al., 2013). After Wnt signaling agonist treatments, the ratios of BrdU positive palatal mesenchymal cells were restored in *Pax9^-/-^* samples when compared to *Pax9^+/-^* tissues (Fig. 7B).

**Figure 7.**
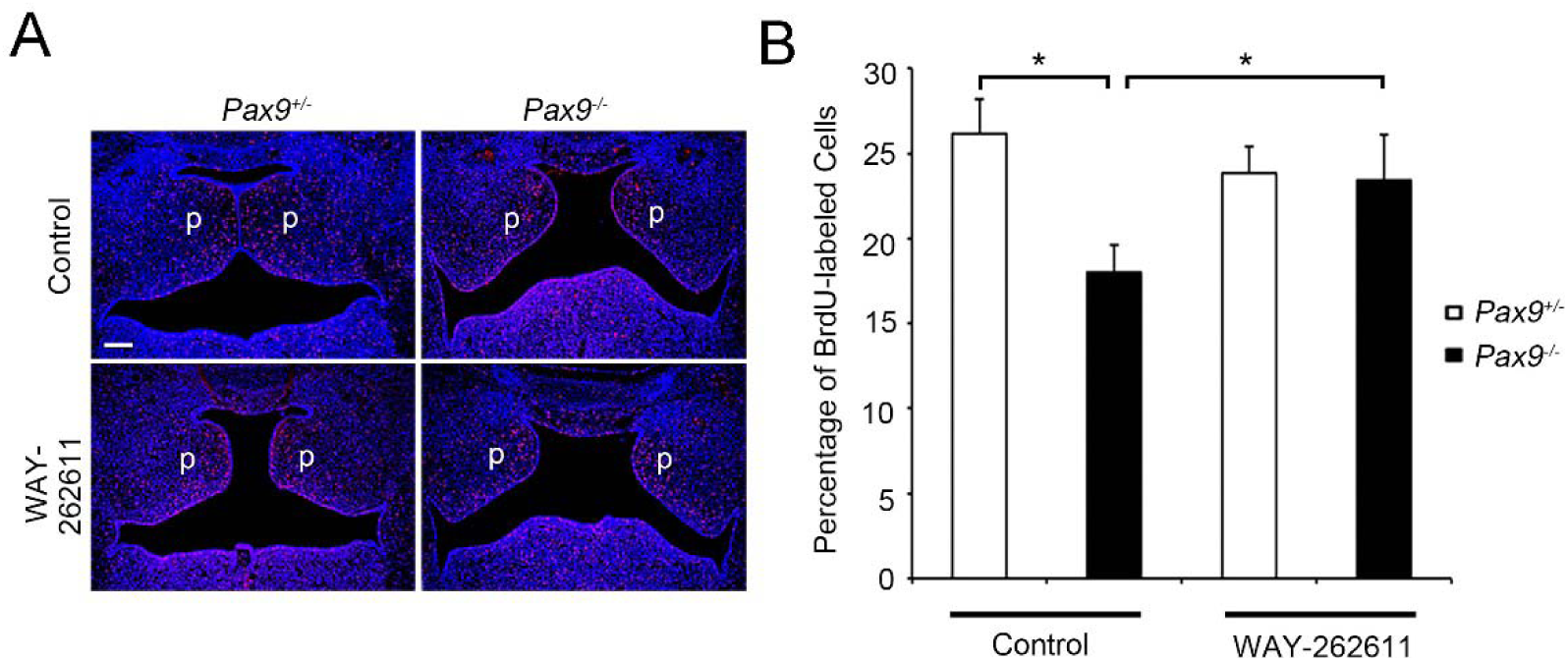
Restored cell proliferation in the palate of WAY-262611 treated *Pax9^-/-^*mice. (A) Representative images of sections through the posterior regions of the palate in E14.5 *Pax9^+/-^* (left column) and *Pax9^-/-^* (right column) embryos without (upper row) or with WAY-262611 treatment (lower row). Red signals showed the distribution of BrdU-labeled nuclei. Blue signals showed the total nuclei, which were stained with Hochest. (B) The percentage of BrdU-labeled cells in the E14.5 palatal mesenchyme. n=5, error bar represents s.d. *, *P*<0.01.

Since Wnts are known to direct the commitment to and differentiation along the osteoblastic lineage (Baron and Kneissel, 2013), we compared alkaline phosphatase activity and type I collagen *(Col1a1)* mRNA expression in treated and untreated *Pax9^+/+^* and *Pax9^-/-^* maxillary and palatine processes. The robust levels of enzymatic activity observed in treated *Pax9^-/-^* tissues closely resembled that in *Pax9^+/+^* tissues and marked the onset of renewed bone formation in these areas (Fig. 8A,B). Alkaline phosphatase is an early osteoblast differentiation marker, which was activated in the osteoprogenitor cells of the palatal mesenchyme at E15.5 in the wildtype control embryos. However, no alkaline phosphatase activity was detected in the palate of *Pax9* mutant embryos. After Wnt agonist treatment, enzymatic activity was restored in the *Pax9* mutant palate (Fig. 8A). *Col1a1,* a marker of differentiated osteoblasts appeared in osteoblasts within osteogenic zones as early as E15.5 and increased from E16.5 to E18.5 in the wildtype embryos. In contrast, *Pax9* mutant samples failed to show *Col1a1* expression in putative mesenchyme. After Wnt agonist treatment, *Col1a1* expression was restored as indicated by the osteogenic activity within the palatal shelves (Fig. 8B).

**Figure 8.**
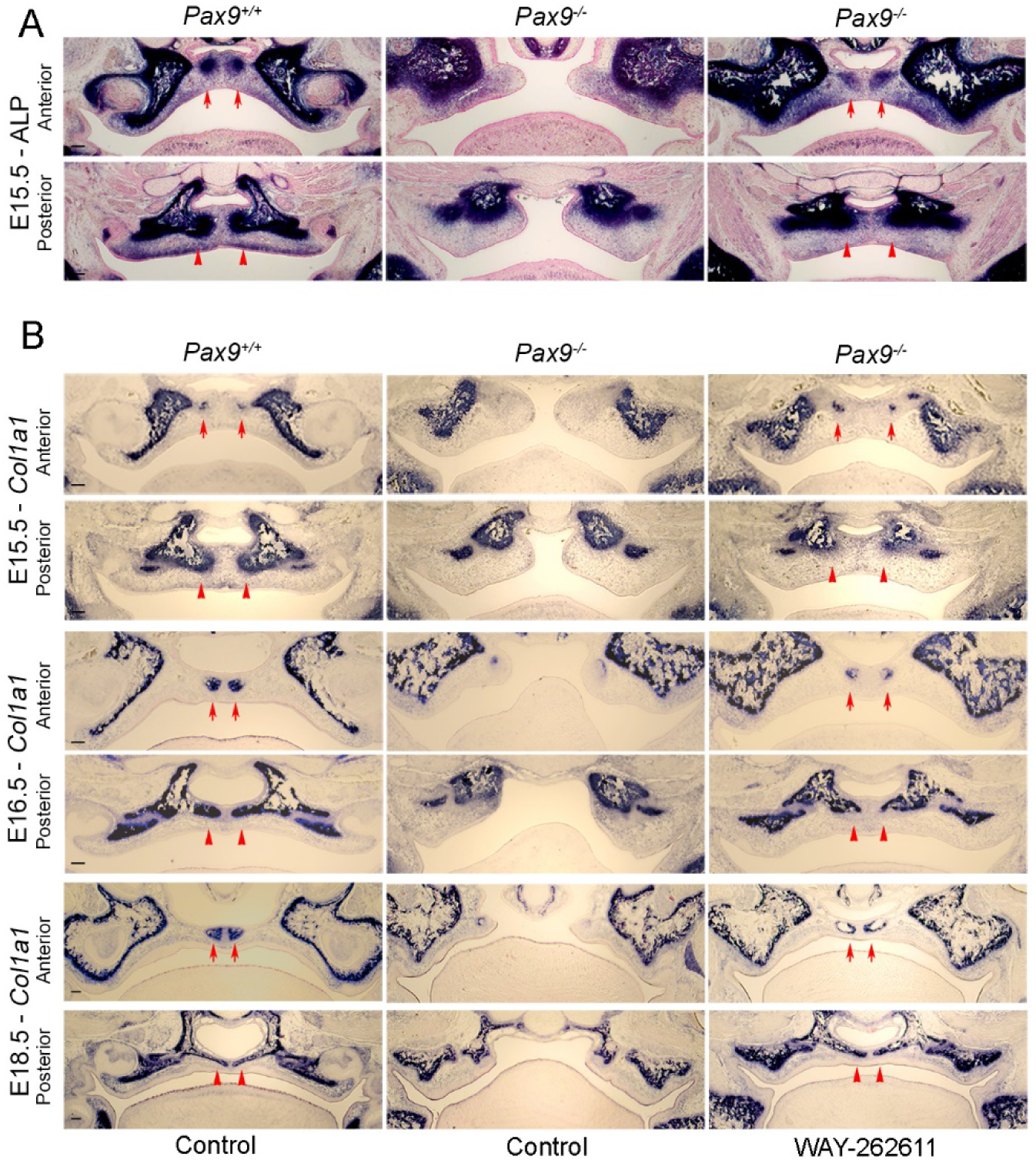
Restored osteogenesis in the palate of WAY-262611 treated *Pax9^-/-^* mice. (A) Detection of alkaline phosphatase expression in the palatal mesenchyme during palatal bone formation in *Pax9^+/-^* and *Pax9^-/-^* embryos without or with WAY-262611 treatment. Position-matched frontal sections of E15.5 embryos through the anterior and posterior palates are used. Alkaline phosphatase staining is shown in blue color. Red arrows, osteogenic centers of the developing palatal processes of the maxilla. Red arrowheads, prospective palatal processes of the palatine bone. (B) Comparison of expression of *COL1a1* mRNA in the palatal mesenchyme during palatal bone formation in *Pax9^+/-^* and *Pax9^-/-^* embryos without or with WAY-262611 treatment. Position-matched frontal sections of E15.5, E16.5 and E18.5 embryos through the anterior and posterior palates are used. mRNA signals are shown in blue color. Red arrows, osteogenic centers of the developing palatal processes of the maxilla. Red arrowheads, prospective palatal processes of the palatine bone. n=5 for each assay.

Taken together, these data establish that the IV delivery of small-molecule Wnt agonists to pregnant *Pax*9^+/-^females within a critical gestational window was effective in bringing about successful closure of the palatal shelves in *Pax9^-/-^* embryos (Fig. 8). While Wnt signaling was restored in treated palates, Wnt therapies did not correct the developmental arrest of tooth organs, thymus, parathyroid glands and hind limb defects in treated mutants (Fig. S4). Neither did the Wnt therapy lead to prolonged survival of treated *Pax9^-/-^* neonates, suggesting that calcium homeostasis remained severely disrupted in the absence of *Pax9,* likely leading to abnormal cardiac function. The MRI imaging showed that *Pax9^+/-^* mothers injected with small-molecule Wnt agonists, as well as their wildtype and heterozygous progeny showed no toxic effects or tumor development up to 18 months after the last tail-vein injection (Data not shown).

## Discussion

These studies explored, for the first time, the nature of the molecular relationship between *Pax9* and Wnt genes in palatogenesis. We provide definitive evidence that *Pax9* deficiency alters Wnt expression activity in palatal shelves. We confirm that Pax9 shares a direct upstream relationship with the Wnt pathway as the transcription factor can directly bind to the promoters of *Dkk1* and *Dkk2,* proteins that inhibit the binding of Wnt ligands to the LRP receptor complex. We confirmed this relationship by delivering small molecule Dkk inhibitors (Wnt agonists) into the tail-veins of pregnant *Pax9^+/-^* mice during a critical developmental window for palatogenesis. Such interventions restored Wnt signaling in palatal mesenchyme and specifically increased cell proliferation within posterior palatal mesenchyme, where *Pax9* and *Dkks* co-localize. Wnt activation in *Pax9^-/-^* palates promoted osteogenesis in the region of the merging shelves, consistent with the role Wnts play in promoting osteoblast differentiation. Collectively, these data demonstrate that the Pax9/Wnt pathway is critical for the formation of the secondary palate.

### Pax9 and the Wnt signaling pathway play key roles in secondary palate formation

How molecular pathways and cellular processes are integrated to coordinate palate morphogenesis is critical to our understanding of the mechanisms that lead to clefts of the secondary palate, the most common among craniofacial disorders. Studies of knockout mice that are characterized by a consistent cleft palate phenotype, have provided valuable insights into how single gene deletions disturb palatogenesis (Greene and Pisano, 2010; Bush and Jiang, 2012; Funato et al., 2015). In addition, gene expression studies point to a remarkable molecular heterogeneity along the anterior-posterior axis that is present at early stages of palatal shelf outgrowth. This morphogenetic gradient is critical for the proper patterning, outgrowth and fusion of the palatal shelves and involves different cascades of gene expression in the anterior, posterior and mediolateral regions of the palatal shelves (Bush and Jiang, 2012). *Msx1* and *Shox2* are examples of genes that are restricted to anterior palatal mesenchyme and whose expression is regulated by Bmp signaling (Li and Ding, 2007). Beads soaked in Bmp4 were able to induce *Msx1* expression in anterior but not posterior palatal mesenchyme, confirming that palate formation involves temporospatially regulated molecular networks of transcription factors and growth factors (Hilliard et al., 2005).

Our studies utilized the *Pax9^-/-^* mouse model to further advance our understanding of the gene networks that regulate palate formation. So far, studies have revealed that *Pax9* is expressed throughout palatal mesenchyme but that its expression is most dominant in the posterior region where it regulates epithelial-mesenchymal signaling interactions and AP patterning. The lack of *Pax9* leads to defects in the outgrowth of palatal shelves that is linked to decreased levels of cell proliferation in the posterior mesenchyme (Zhou et al., 2013). In contrast, *Osr2* that is downstream of *Pax9* is also expressed throughout palatal mesenchyme but specifically affects cell proliferation in the lateral region (Lan et al., 2004). Hence, there is discrete molecular control of Pax9’s actions in posterior palatal mesenchyme that remains to be identified.

Our unbiased studies of gene expression profiles in *Pax9^-/-^* palatal shelves revealed that the expression of Wnt signaling pathway genes are altered in the absence of *Pax9.* Wnt proteins are highly conserved secreted signaling molecules that regulate key developmental processes. At least 12 of the 19 known Wnt ligands are expressed in murine palates (He et al., 2008; Warner et al., 2009) and *WNT5a* mutations have been associated with human cleft palate (Yamaguchi et al., 1999). The expression of *Dkk1* and *Dkk2,* inhibitors of Wnt ligand binding to the LRP receptor complex, were markedly altered in *Pax9^-/-^* palates. Hence, we analyzed how Pax9 affects the canonical Wnt signaling pathway through its control of *Dkk* expression.

### The exclusive correction of Pax9-mediated cleft palate defects with Wnt agonists underscore the importance of this molecular pathway in secondary palate formation

The surprising fidelity with which palatal defects are corrected in *Pax9^-/-^* embryos, through the controlled intravenous delivery of Wnt agonists can be explained by the fact that Wnt signaling genes are potent downstream effectors of Pax9’s functions in palatal mesenchyme. Interestingly, such interventions could not restore the development of the thymus, parathyroid glands, and ultimobranchial bodies or correct other defects such as supernumerary preaxial digits. It is hence likely that the direct molecular relationship between *Pax9* and the Wnt genes during palate formation is not shared by other affected organs. Hence, organs, other than the secondary palate, that are derived from the pharyngeal pouches may require essential contributions from downstream effector pathways such as Bmp, Fgf and Shh. It is also likely that Wnt agonist delivery and dosage schedules that proved optimal for palatogenesis would have to be modified to achieve the correction of other organ defects in *Pax9^-/-^* mice.

The interventions with small molecule Wnt agonists failed to prolong the survival of *Pax9^-/-^* neonate pups from treated pregnant dams as they succumb at birth. While earlier studies have noted that *Pax9^-/-^* newborn pups die shortly at birth, most likely as a consequence of secondary palate clefts (Peters et al., 1998), our data suggest that systemic alterations may lead to early death. These can be attributed to a severe disruption in calcium homeostasis that may compromise cardiac function as supported by lethality of *Pax9^-/-^* newborns with palatal defects that were corrected *in utero.*

Like cleft secondary palates, the failure of tooth organs to advance beyond the bud stage is a consistent phenotypic feature seen in *Pax9^-/-^* mice (Peters et al., 1998; Zhou et al., 2011). It is intriguing that Wnt agonists when delivered intravenously during a critical developmental window for odontogenesis, only advanced tooth organs to the early cap stage. In humans, mutations in *PAX9* are mainly associated with congenitally missing posterior teeth (Goldenberg et al., 2000; Stockton et al., 2000), thus underscoring its key role in the formation of patterned dentition. *Pax9* has been shown to play a critical role in murine dental mesenchyme by influencing epithelial-mesenchymal signaling events that drive tooth morphogenesis (Peters et al., 1998; Nakatomi et al., 2010). While the canonical Wnt signaling pathway has emerged as a key driver of tooth signaling interactions and is likely downstream of *Pax9* in dental mesenchyme (O'Connell et al., 2012), the precise nature of their molecular relationship during odontogenesis is not known. *Pax9* also serves as an important integrator of mesenchymal signaling events that drive tooth morphogenesis and that involve *Bmp4, Fgf3* or *Fgf10* (Peters et al., 1998; Nakatomi et al., 2010). Hence, combinatorial therapies involving Wnt agonists and these growth factors may prove more effective in advancing tooth morphogeneis and should be the focus of future studies.

We also observed that the intravenous delivery of Wnt agonist small molecules failed to rescue cleft defects in 50 *Msx1^-/-^* mice exposed to Wnt agonists. *Msx1* is a partner gene of *Pax9* that also plays a role in palatogenesis and is more dominantly expressed in the anterior zone of palatal mesenchyme. In contrast, *Pax9* and the *Dkks* are more restricted to the posterior region. Hence, a morphogenetic gradient of differential gene expression along the anterior-posterior axis controls the patterning and closure of palatal shelves (Hilliard et al., 2005; Zhou et al., 2013). It is hence likely that downstream effector genes other than the Wnts mediate *Msx1’s* actions in anterior palatal mesenchyme.

### Clinical implications for the use of small molecule replacement therapies for the treatment of developmental disorders of the craniofacial complex

The wide implications of the Wnt-ß-catenin signaling pathway in many developmental processes and in adult tissue homeostasis has encouraged development of pharmacological modulators of this pathway (Clevers and Nusse, 2012; Baron and Kneissel, 2013; Kahn, 2014). Previously, a small-molecule-based chemical genetic approach restored the stability and function of GSK-3ß in *GSK-3ß^FRB^*^/FRB^\** mice, such that after rapamycin treatment, cleft defects were rescued in 6 of 9 mutant pups (Liu et al., 2007). New therapies that block the function of the Wnt antagonist, *sclerostin,* restored bone mass and strength in patients at risk for fractures. Individuals receiving antibodies to sclerostin show a decrease in bone resorption markers, thus underscoring the safety and efficacy of such approaches for the treatment of human osteoporosis (Recker et al., 2015). Several preclinical studies also report that other therapies targeting Dkk1 inhibition induce bone gain (Baron and Kneissel, 2013).

Our data indicate that the intravenous administration of small molecule Dkk inhibitors led to an increase in Wnt signaling in palatal mesenchyme during embryonic development. This resulted in an increase in cell proliferation and osteoblast differentiation. While the key effects of Wnt agonists such as WAY-262611 and IIIc3a for the closure of palatal clefts can be explained on the basis of their effects on palatal osteogenesis, questions naturally arise about how such approaches can be translated into therapies for human cleft palate disorders. Since Wnt signaling is key to the development of several organs, activation of the pathway through exogenous Wnt agonist therapies could lead to abnormalities in other organ systems. Our histopathologic and MRI imaging analyses of pregnant *Pax9^+/-^*females that received Wnt agonist injections and surviving *Pax9^+/+^* or *Pax9^+/-^* littermates showed no toxic effects or tumor development up to 18 months following the last tail-vein injection. Hence, small doses of Wnt agonists that target inhibitors of receptor binding delivered in a controlled manner during a specific developmental window only affect cells like those in the palatal mesenchyme that have not reached threshold levels needed for the hierarchy of Wnt signaling.

Although ultrasound technologies can diagnose cleft palate conditions as early as 13 weeks of gestation, small-molecule therapies delivered in low doses to pregnant mothers are likely to be fraught with difficulties and will require further experimentation on risk to benefit ratios. Alternate strategies to reduce exposure of the mother could include delivery into the amniotic fluid as shown to be effective for the delivery of recombinant ectodysplasin (Hermes et al., 2014). Alternatively, early postnatal interventions that utilize the well-timed local delivery of Wnt agonist small-molecules or proteins during surgical correction procedures may prove effective in bringing about the timely closure of palatal shelves.

## Materials and Methods

### Mouse Strains

All animal procedures were approved by the Institutional Animal Care and Use Committee (IACUC) at the University of Utah (Protocol #13-12010). *Pax9^+/-^* and *Msx1^+/-^* mice were provided by Dr. Rulang Jiang (Cincinnati Children’s Hospital) (Zhou et al., 2011) and Dr. Yiping Chen (Tulane University) (Satokata and Maas, 1994) respectively. *BAT-Gal* mice for Wnt activity (Maretto et al., 2003) (JAX 005317 from Jackson Laboratory). Whole heads of E13.5 and E14.5 *Dkk1^Tg107-LacZ^* embryos were provided by Dr. Ulrich Rüther (Heinrich-Heine-University) (Lieven et al., 2010). The *Pax9^+/-^, Msx1^+/-^* and *BAT-Gal* mice were maintained in *C57BL/6* background, 2-8 months old females were used for intercross mating, mice for testing toxic effects were 18 months old.

### Gene expression analysis

Whole transcriptome profiling of the developing palatal shelves were analyzed by RNA-Seq as previously described (Jia et al., 2013). In brief, total RNAs were extracted from individual palatal shelves using the RNeasy Micro Kit (Qiagen). cDNA templates were generated using the Illumina TruSeq RNA Sample Prep Kit v2 with oligo(dT) selection. Sequenced reads were mapped to the reference mouse genome (mm 10) using Novoalign (v2.08.03). Read counts were generated using USeq’s Defined Region Differential Seq application and normalized counts were used in DESeq2 to measure differential expression. The RNA-seq raw data have been deposited into Gene Expression Omnibus database (GEO) (http://www.ncbi.nlm.nih.gov/geo) under accession number GSE89603. The candidate genes were screened from a comparison of 5 sets of data through the procedure below: First, the normalized counts were cut-off at 10 in at least one of the samples, then the fold change was chosen at 1.5-fold or higher and at P-values <0.01 from the Audic Claverie test with Benjamini Hochberg FDR multiple testing correction. Next, the gene list was sorted by p value and lists of genes were submitted to DAVAD (https://david.ncifcrf.gov) or Toppgene (https://toppgene.cchmc.org) for functional enrichment analysis.

### Real-time RT-PCR

Palatal shelves were micro-dissected from E13.5 and E14.5 embryos. After genotyping, total RNA was extracted individually using the RNeasy Micro Kit (Qiagen). First-strand cDNA was synthesized using the SuperScript First-Strand Synthesis System (Invitrogen). Quantitative PCRs were performed in a StepOnePlus™ Real-Time PCR System (Applied Biosystems) using the SYBR Green^ER^ qPCR Supermix (Invitrogen). Gene-specific primers are listed in the supplemental data. For each sample, the relative levels of target mRNAs were normalized to HPRT using the standard curve method. 5 sets of samples were analyzed for each gene. Student’s t-test was used to analyze differences and P values <0.05 considered statistically significant.

### Delivery of Small-Molecule Wnt Pathway agonists

Two small molecules agonists were used to treat the timed pregnant female mice. WAY-262611, (1-(4-(Naphthalen-2-yl)pyrimidin-2-yl)piperidin-4-yl)methanamine, is a 2- aminopyrimidine compound (Fig. S1A) that antagonizes the effect of Dkk-1 by preventing the formation of the Dkk1/LRP5/Kr2 complex with 6-8 hours half-life (*t*_1/2_) in plasma (Pelletier et al., 2009). It inhibits Dkk1 at EC_50_=0.63 μΜ from TCF-luciferase assay. IIIc3a, 9-Carboxy-3-(dimethyliminio)-6,7-dihydroxy-10-methyl-3H-phenoxazin-10- iumiodide, is an enhanced in-solution stable gallocyanine analog (Fig. S1B), disrupts the interaction of LRP5/6 with Dkk (Dickkopf) in a competitive manner. EC_50_=5 μΜ in LEF-luciferase assay using NIH3T3 cells (Li et al., 2012).

WAY-262611 and IIIc3a (Millipore, Billerica, MA, USA) were dissolved 25 mg/ml in DMSO. Stock solutions were diluted in PBS and counted at 12.5 or 25 mg/kg per mouse body weight prior to injection into the tail vein as previously described (Kowalczyk et al., 2011). Control treatments consisted of tail-vein injections of 10% DMSO in PBS. Mice were monitored for any discomfort or side effects.

### Histology and *In-situ* hybridizations

Embryos were fixed in 4% paraformaldehyde (PFA) overnight, the 7 μm-thick paraffin sections stained with H&E for microscopic evaluation. In addition, serial coronal sections were stained with Massons trichrome (Chen et al., 2009) to evaluate connective tissue deposition. In-situ hybridizations were performed using digoxigenin-labeled RNA probes to *Dkk1, Dkk2* and *Col1a1* as described previously (D'Souza et al., 1993; Zhang et al., 1999). 1 μg/ml antisense RNA probe was loaded on each section and anti-digoxigenin-AP antibody (11093274910, ROCHE, 1:1000) was used to detect the labeled probe. Comparable images were taken by digital microscope (EVOS). For each assay, at least 5 replicates were performed.

### Gross examinations, histopathologic and MRI analyses

All mouse embryos, neonates and adults were first evaluated through full-body visual examination. Whole embryos were fixed in 10% neutral buffered formalin, 5 μm-thick transverse paraffin sections were stained with H&E for microscopic evaluation. The adult mice were anaesthetized with isoflurane and fixed by whole-body perfusion. The magnetic resonance imaging (MRI) was performed using 7 Tesla Bruker BioSpec MRI scanner under the following parameters: A T2 weighted RARE scan was acquired using a 7.2 cm-diameter quadrature radiofrequency transmitter-receiver (Bruker Biospec), with 125 μm isotropic resolution, echo train length of 4, matrix size of 768x256x256 and TE/TR = 41/1200 ms on a Bruker Biospec 70/30 instrument (Bruker Biospin, Ettlingen, Germany).

### Whole mount LacZ staining and alkaline phosphatase staining

For whole mount LacZ staining, the embryos were fixed in 1% PFA then treated as previously described (Lan et al., 2004). For alkaline phosphatase (ALP) staining, embryos were fixed in 4% paraformaldehyde overnight. The frozen sections were incubated in the NTMT buffer (100 mM NaCl, 100 mM Tris-HCl pH 9.5, 50 mM MgCl2, 0.1% Tween-20) containing 4.5 μl/ml nitroblue tetrazolium (NBT) and 3.5 μl/ml 5-bromo-4-chloro-3-indolyl phosphate (BCIP) for 20 min RT (Baek et al., 2011). Comparable images were taken with a digital microscope (EVOS) or a stereomicroscope (Zeiss Stemi 508). For each assay, at least 5 replicates were used to establish reproducibility of results.

### BrdU labeling and cell proliferation assay

Timed pregnant female mice were injected once intraperitoneally at E14 with the BrdU Labeling Reagent (Roche, 15 μl/g body weight). Embryos were harvested two hours later. The 5 μm paraffin sections were rehydrated and submerged in acetone. After proteinase K digestion, HCl denature, blocking solution (2% BSA, 10% goat serum, 0.1% Tween in PBS) incubation, samples were incubated with Alexa Fluor 594- conjugated anti-BrdU antibody (Invitrogen, #B35132) solution (1:50 in blocking solution) overnight. After counterstain with Hoechst, images were obtained using a Zeiss fluorescence microscope and analyzed with Imaris software (Zhou et al., 2013). The cell proliferation data were recorded and analyzed from 5 independent control and mutant littermate pairs. Student’s t-test was used to analyze the differences in the datasets and P <0.01 was considered statistically significant.

### ChIP-qPCR assay

The plasmid pCMV-Tag3B-Pax9 was transfected into HEK293T cells (From ATCC). ChIP was performed as previously described with modification (Park et al., 2012). Briefly, cells were cross-linked with 1% PFA. After quenching in glycine and rinsing in PBS, cells were re-suspended in 1ml lysis buffer followed by sonication. A 50 μl aliquot was saved as input, and the rest was incubated overnight with Dynal Protein G magnetic beads (Thermo Fisher, 10003D) that had been pre-incubated with normal IgG (Covance, # MMS-126R) or anti-Myc (EMD Millipore, 05-724) antibody. After washed with RIPA buffer, LiCl buffer and TE buffer, bound complexes were eluted by heating at 65°C and crosslinking was reversed by overnight incubation at 65°C. After purified by Agencourt AMPure XP beads (Beckman Coulter, A63880), the concentration was determined by Qubit 3.0 Fluorometer (Thermo Fisher). 0.2 ng of each DNA sample was used for qPCR analysis. The primer sequences for *Dkk1* and *Dkk2* promoter regions that were used in the qPCR are listed in the supplemental data.

### Whole palate shelf organ culture

The WAY-262611 treated palates were dissected on ice by removing the lower jaw and brain, samples with residual fusion defects were marked and cultured as previously described (Almaidhan et al., 2014). *Pax9^+/-^* and untreated *Pax9^-/-^* embryos were cultured as control. The dissected E18.5 palate shelves were put into 2 ml of medium (CO2 independent medium, Invitrogen, #18045-088; 20% FBS; 1X antimycotic-antibiotic, Invitrogen, #15240-096). Culture tubes were rotated 12 rpm at an angle of 20 degree in a 37°C incubator for 3 days with daily media changes. After pictures taken with a stereomicroscope, tissues were fixed and embedded for histologic analyses. At least 5 replicates of each treatment and genotype were utilized for culture experiments.

## Acknowledgments

We thank Dr. Rulang Jiang and Dr. YiPing Chen for donating breeding pairs of *Pax9* and *Msx1* heterozygote mice, respectively, and for their advice throughout the project. We thank Dr. Ulrich Rüther for providing fixed heads of *Dkk1^Tg107-LacZ^* samples. We are also grateful to Dr. Mengyao Tan for her guidance in working out the conditions for the ChIP-qPCR assays and to Dr. Jeffrey Pelletier for sharing his insights on properties of WAY-262611. The technical assistance of Mr. Greg Pratt is also acknowledged.

## Author contributions

S.J. designed and performed RNASeq and hybridization experiments, mouse tail-vein injections and breeding. He also carried out data analysis, prepared the figures and contributed to manuscript writing; J.Z. performed ChIP and qPCR assays, BrdU staining and counts as well as whole-mount hybridizations; C.F. prepared histologic sections and assisted with animal injections, alkaline phosphatase staining and in-situ hybridizations as part of his dental student summer research project; Y.W. prepared the samples for MRI and assisted with qPCR assays and BrdU positive cell counting; J.B. performed the initial round of small-molecule Wnt agonist therapy as part of his Ph.D. dissertation project; P.S. provided crucial guidance in the design of these experiments and contributed comments on the manuscript along with G.M. who helped conceive the study with R.D.S. All experiments were designed and data analyzed by R.D.S. who wrote the manuscript.

## Data availability

The RNA-seq raw data have been deposited into NCBI Gene Expression Omnibus database (GEO) (http://www.ncbi.nlm.nih.gov/geo) under accession number GSE89603. All material requests and correspondence should be addressed to R.D.S.

